## Supplementary Information for "Multi-condensate state as a functional strategy to optimize the cell signaling output"

### Supplementary Texts

#### Supplementary Text 1: Surface density of Arp2/3 in the n-cluster configuration for perfect hemisphere (hemiSphericity = 1)

Suppose we have a large hemispherical cluster of radius R. We can split the large cluster into n smaller ones, each having a radius of r. Total number of Arp2/3 is A.

Volume (mass) conservation reads: $\frac{2}{3}\pi R^3 = n * \frac{2}{3}\pi r^3 \Rightarrow R = n^{\frac{1}{3}} r$

Surface area of the large cluster, $S = 4\pi R^2$

Surface area of a smaller cluster, $s = 4\pi r^2 = 4\pi R^2 * n^{-\frac{2}{3}} = S * n^{-\frac{2}{3}}$

Surface density of Arp2/3 at the large cluster, $\alpha = \frac{A}{S}$

Surface density of Arp2/3 at a smaller cluster, $\rho = \frac{\frac{A}{n}}{s} = \left(\frac{A}{S}\right) * n^{-\frac{1}{3}} = \alpha * n^{-\frac{1}{3}}$

#### Supplementary Text 2: Derivation of Arp2/3 threshold density (single cluster case)

For one cluster case,

$$\frac{dF}{dt} = k1 * G^2 * \alpha * F - k2 * F \text{ where } G + F = C \text{ (constant)}$$

$$f = dF/dt = k1 * (C - F)^2 * \alpha * F - k2 * F$$

After differentiating w.r.t. F,

$$f' = (\alpha k1C^2 - k2) - 4\alpha k1C * F + 3\alpha k1 * F^2$$

$$f'(F = 0) = \alpha_{critical}\, k1C^2 - k2 = 0 \Rightarrow \alpha_{critical} = \frac{k2}{k1 * C^2}$$

#### Supplementary Text 3: Surface density of Arp2/3 in the n-cluster configuration for oblate hemispheroids (hemiSphericity < 1)

Let's consider a hemispheroidal cluster (Supplementary Fig. 9) with basal radius $r$ and height $h$. When $h < r$, these objects are called oblate spheroids. For a perfect hemisphere, $h = r$.

Now, let's say, to create an oblate hemispheroid from the perfect hemisphere, we multiply the height by a factor $f$, where $0 < f < 1$, such that, $h_o = f * h$

To have volume conservation,

$$r_o{}^2 h_o = r^2 h \Rightarrow r_o = r/\sqrt{f} \quad (3.1)$$

The oblate cluster will have a hemiSphericity,

$$hS_o = h_o/r_o = (f * h)/\left(r/\sqrt{f}\right) = f^{3/2} * hS = f^{3/2}(since\ hS = 1) \quad (3.2)$$

Now, surface area of an oblate hemispheroid (height = h, basal radius = r),

$$S = 0.5 * (2\pi r^2 + \pi(h^2/e)\ log\ ((1 + e)/(1 - e))) \qquad (3.3)$$

Where the ellipticity parameter, $e = \sqrt{(1 - h^2/r^2)} \qquad (3.4)$

First, we want to know the change in surface area when we create a hemispheroid (with an arbitrary f) from a perfect hemisphere with identical volume.

From equations 3.2 and 3.4,

$$e = \sqrt{(1 - (hS)^2} = \sqrt{(1 - f^3)} \qquad (3.5)$$

Combining equations 3.1 and 3.3,

$$S_{oblate} = 0.5 * (2\pi r_o^2 + \pi({h_o}^2/e)\ log\ ((1 + e)/(1 - e)))$$

$$S_{oblate} = 0.5 * (2\pi(r^2/f) + (\pi/e)(r^2 f^2)\ *\ K(e)\ )$$

$$S_{oblate} = 2\pi r^2[0.5 * (1/f + (f^2/2e)\ *\ K(e))]$$

$$S_{oblate} = S_{hemiSphere}\ K(f) \qquad (3.6)$$

Where $K(f)$ is a function of $f$, which determines the surface-to-volume ratio in converting a perfect hemisphere to an oblate hemispheroid. Since the surface-to-volume ratio is higher for a spheroid, the surface density of Arp2/3 ($\alpha$) will decrease as a function of f.

For a fixed number of Arp2/3,

$$S_{oblate} * \alpha_{oblate} = S_{hemiSpehere} * \alpha_{hemiSphere}$$

$$\alpha_{oblate} = \left(1/K(f)\right)\alpha_{hemiSphere} \qquad (3.7)$$

#### Supplementary Text 4: Derivation of local production of F-actin in the n-cluster configuration

First let's consider n = 1 case,

$$\frac{dF}{dt} = k1 * G^2 * \alpha * F - k2 * F \text{ where } G + F = C \text{ (constant)}$$

$$\text{Steady state, } \frac{dF}{dt} = 0 => k1 * (C - F)^2 * \alpha - k2 = 0 \text{ since } F > 0$$

$$F = C - \beta \quad (4.1) \qquad \text{where } \beta = \sqrt{\frac{1}{\alpha} * \left(\frac{k2}{k1}\right)}$$

Now for the n-cluster state,

$$\frac{dF_{local}}{dt} = k1 * G_{local}^2 * \rho(n) * F_{local} - k2 * F_{local} \text{ where } G_{local} + F_{local} = C \ (constant)$$

From Supplementary Text 1, $\rho = \alpha * n^{-1/3}$ for hemispherical clusters.

Following the same scheme as in Equation 3.1,

$$F_{local} = C - \beta * n^{1/6} \qquad (4.2)$$

From equation 3.6, it follows that if the f-parameter (which in turn decides the hemiSphericity) remains the same for a large cluster and a smaller cluster, splitting a large cluster into n smaller ones follows the same scaling laws as described in Supplementary Text 1.

From Equation 4.2, we can write for an oblate cluster,

$$F_{local} = C - \beta' n^{1/6} \qquad (4.3)$$

Where

$$\beta' = \sqrt{(1/\alpha_{oblate})(k2/k1)} = K(f)^{1/2}\,\beta \qquad (4.4)$$

Finally, we have an expression for local production of F-actin near a hemispheroidal cluster with an arbitrary shape parameter ($f$):

$$F_{local} = C - \beta * K(f)^{1/2} * n^{1/6} \qquad (4.5)$$

We note that, $f = 1 \Rightarrow e = 0$. This leads to 0/0 form within the $K(f)$ expression. By applying the limit, $f \rightarrow 1$, and by applying L'Hospital's rule, $K(f) = 1$.

#### Supplementary Figures

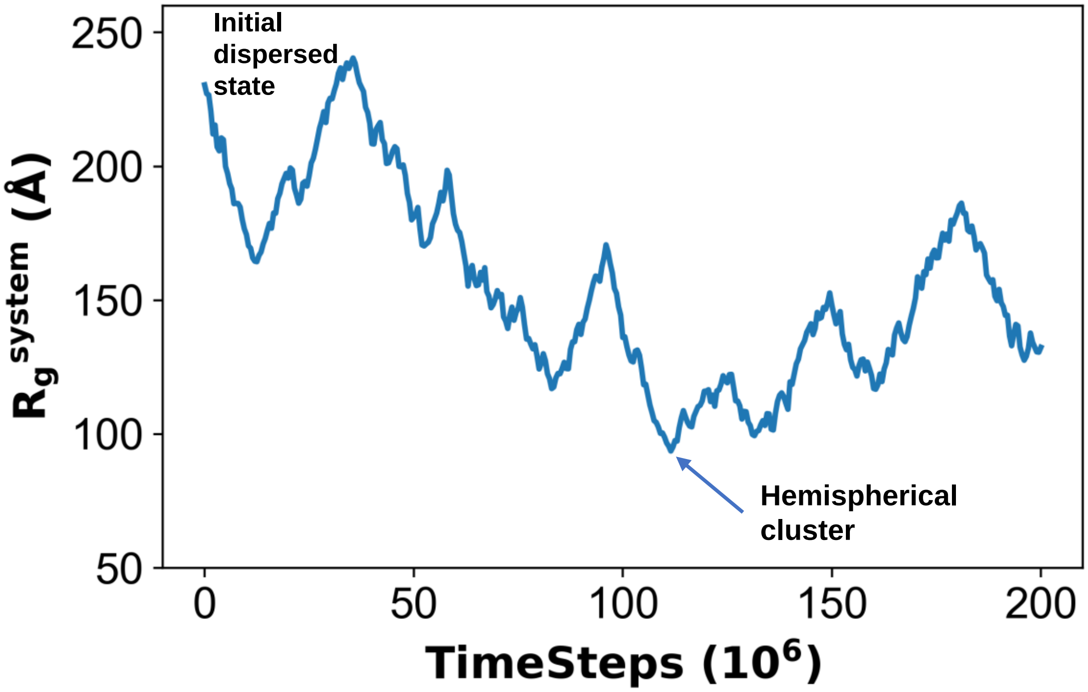


**Supplementary Figure 1: Timecourse of the metadynamics order parameter, $R_g^{system}$.** The initial value indicates the dispersed state while the minima is associated with the hemispherical cluster with minimum surface area.

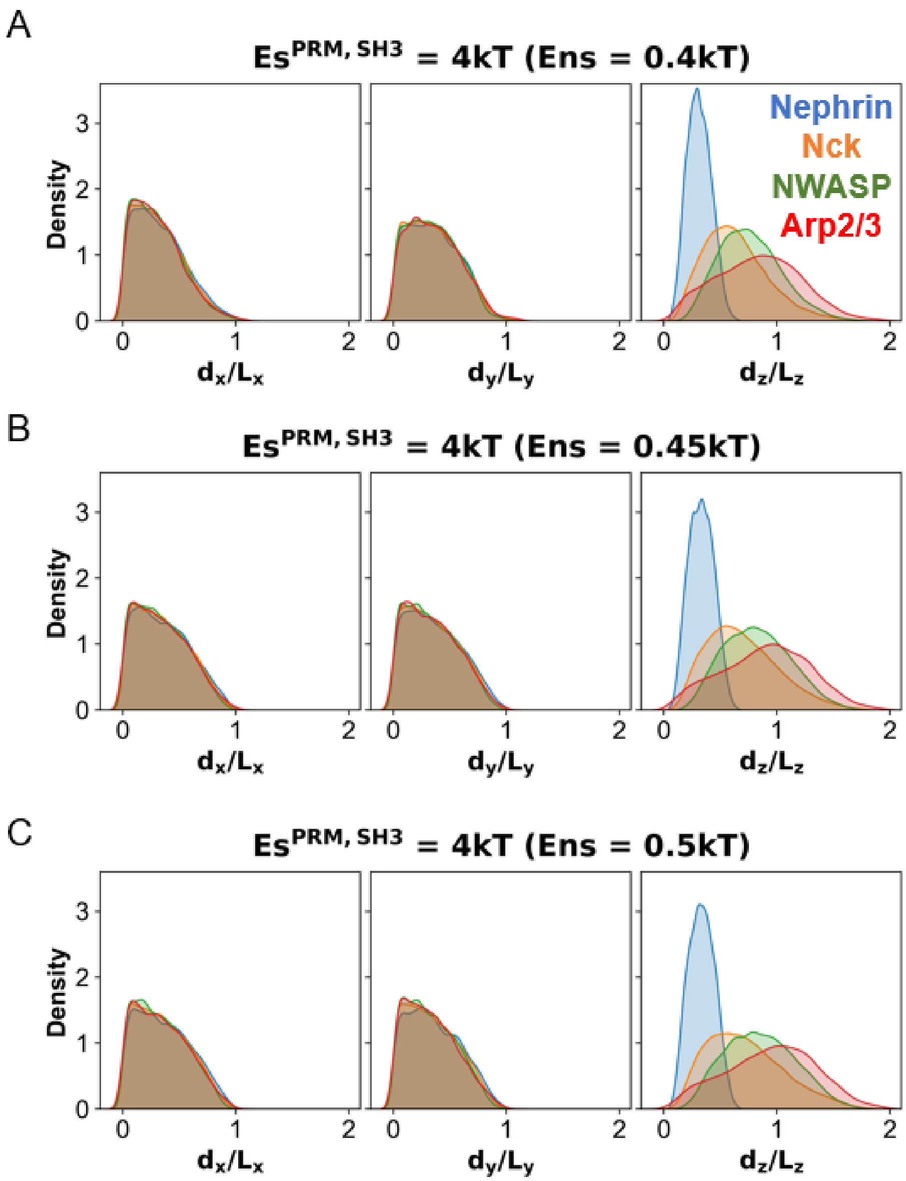


**Supplementary Figure 2: Radial location of different molecular types**, at (A) Ens = 0.4kT (B) Ens = 0.45kT and (C) Ens = 0.5kT. As described in Figure 1F, normalized distances are shown along the X,Y and Z directions.

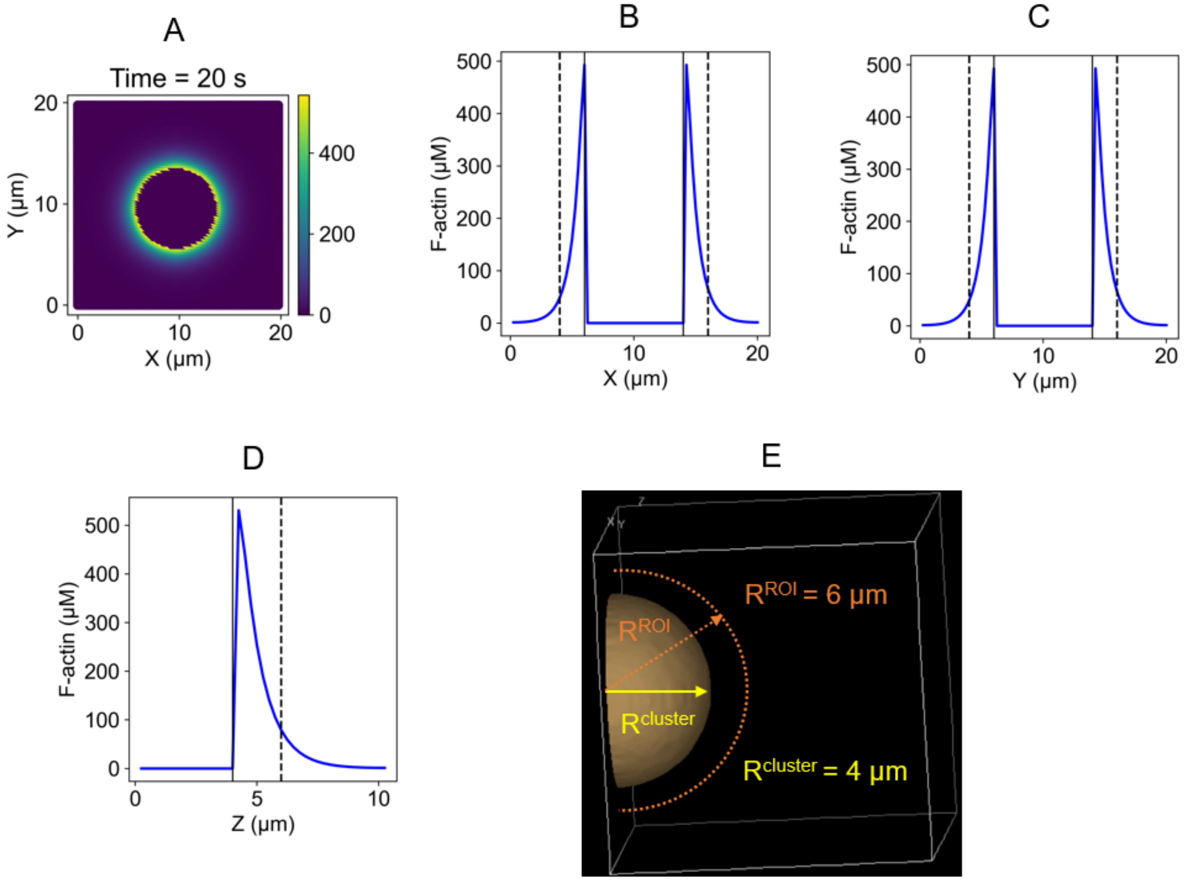


**Supplementary Figure 3: Computation of local F-actin concentration.** (A) Concentration profile of F-actin (FA) at the steady state, as described in Figures 2B and 2C. The basal plane of the condensate, at Z = 0, is displayed. The crossing point of the dotted lines indicate the center (10, 10, 0) of the hemispherical condensate. To compute the local FA concentration near the condensate surface, a hemispherical shell outside the condensate is considered. Along the X,Y and Z directions, concentration profiles are shown (B, C, D). The region between the solid and dashed black lines indicate the local volume which is considered for computing the surface FA concentration. (E) Illustration of the ROI (region of interest) around the hemispherical cluster.

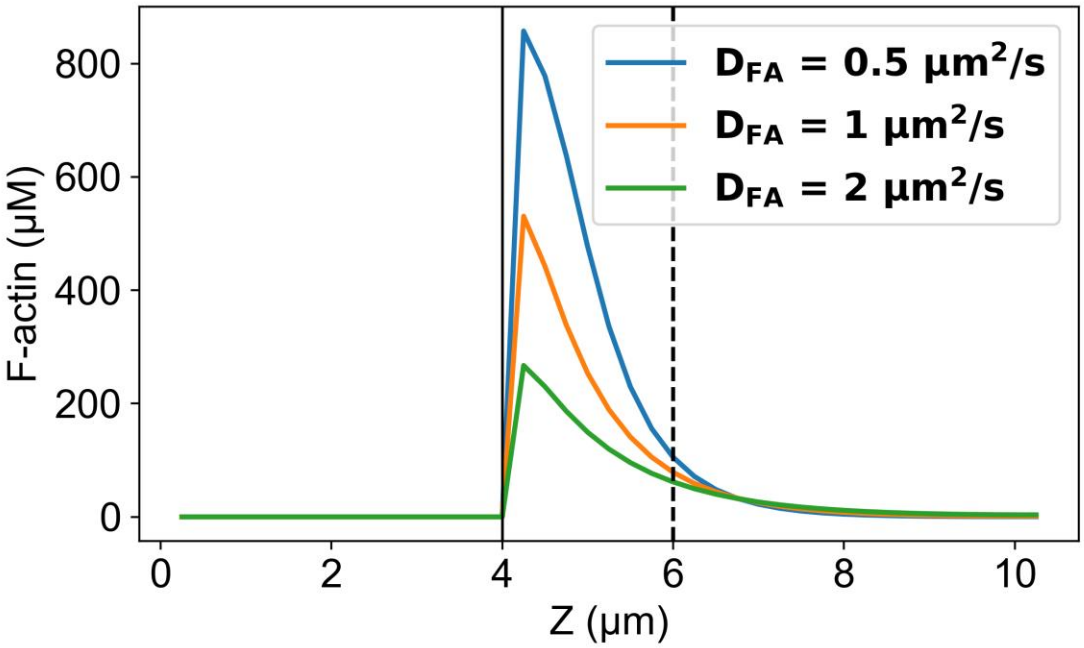


**Supplementary Figure 4: Concentration profiles of F-actin along the Z direction at three different diffusion constants.**

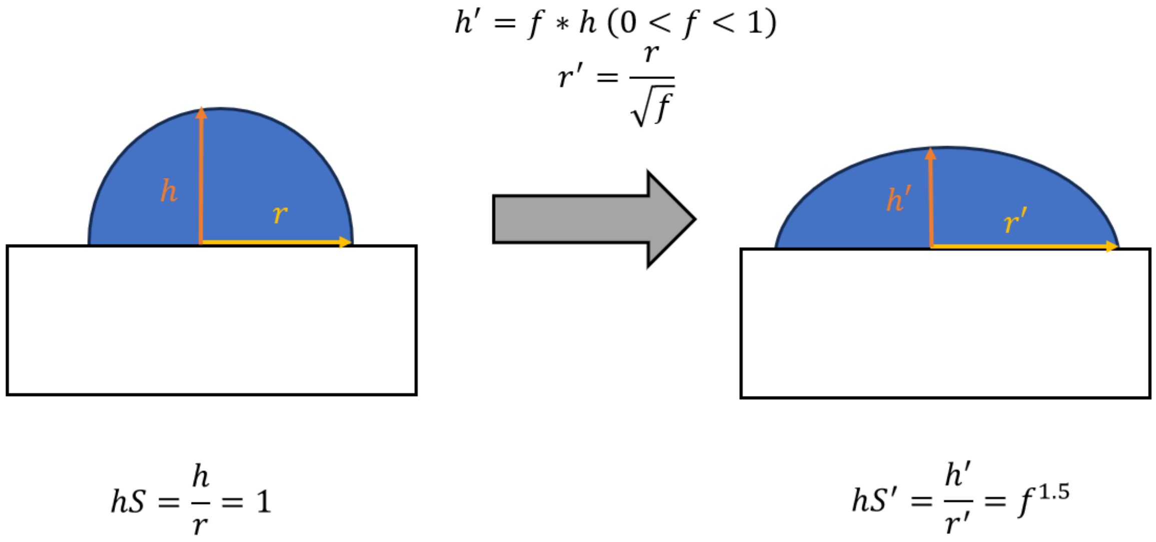


**Supplementary Figure 5: Creating a hemispheroidal geometry from a perfect hemisphere.** For a hemisphere, h = r. HemiSphericity, hS = 1. If we reduce the height by a factor $f$, to converse volume, the radius needs to be increased by a factor of $1/\sqrt{f}$. HemiSphericity of the resultant object will scale with $f^{1.5}$.

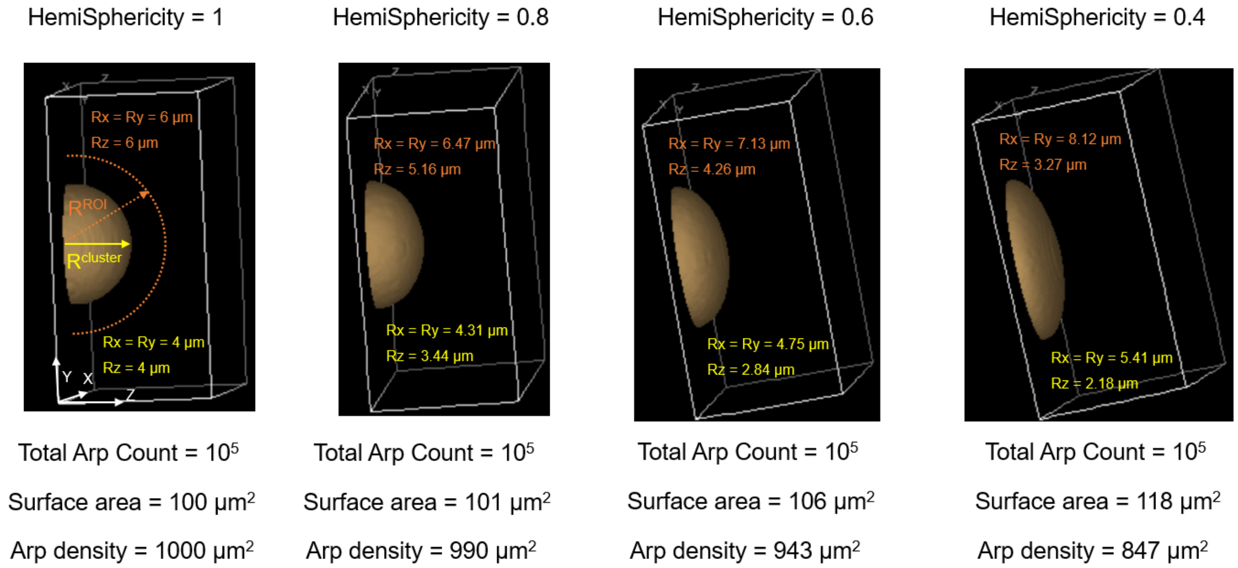


**Supplementary Figure 6: Creating clusters with different degree of hemiSphericity.** With lowering of hemiSphericity, surface area increases. To keep the Arp2/3 count fixed across these scenarios, surface density of Arp2/3 is adjusted. The regions of interest (ROIs) follow the same shape as the cluster in such a way that the local volume within the hemispherical shell remains roughly the same. Color code, yellow: cluster, orange: ROI

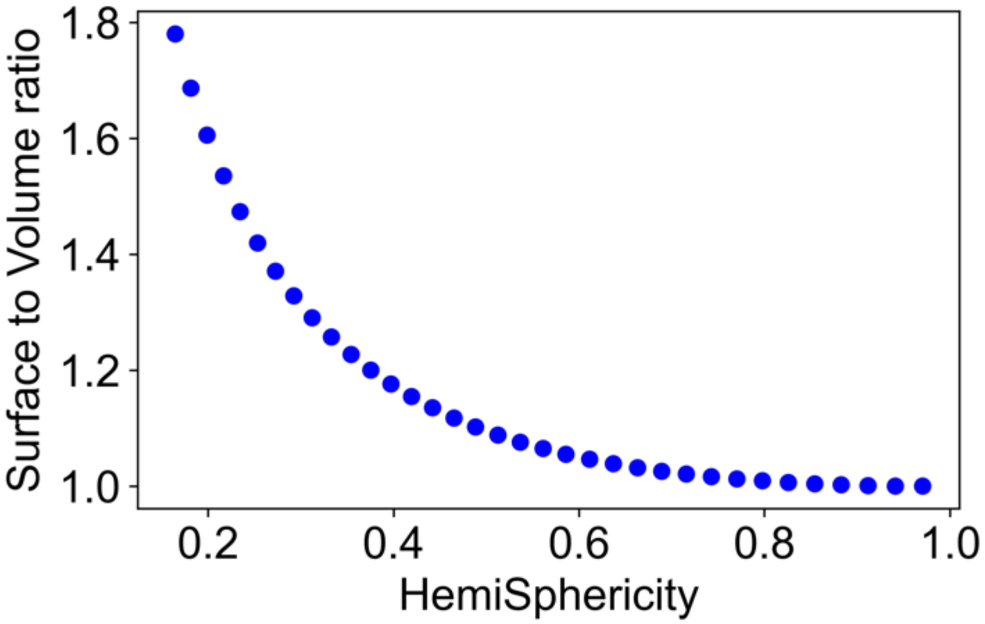


**Supplementary Figure 7: Trend of surface to volume ratio as a function of hemiSphericity.** Keeping volume constant, creating oblate spheroids from a sphere causes a gradual increase in the surface area.

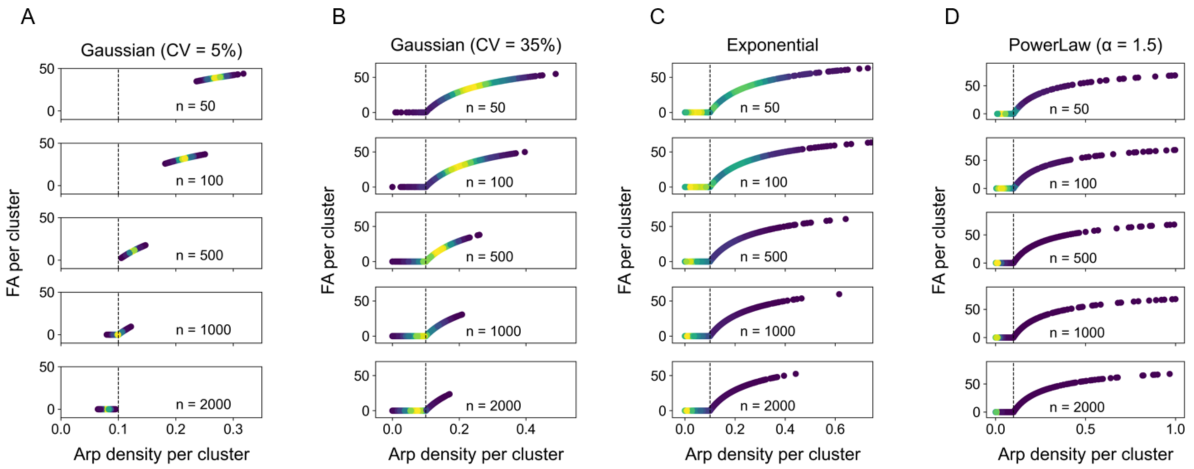


**Supplementary Figure 8: Local F-actin production profiles for different cluster size distributions.** Cluster size follows (A) Gaussian distributions with CV = 5%, (B) Gaussian distributions with CV = 35%, (C) Exponential distributions, (D) Power Law distribution with a shape parameter (α) = 1.5. The cluster numbers (n) are indicated for each panel. The color code indicates the frequency. Darker (purple) ones are less frequent while brighter (yellow) ones are more frequent.

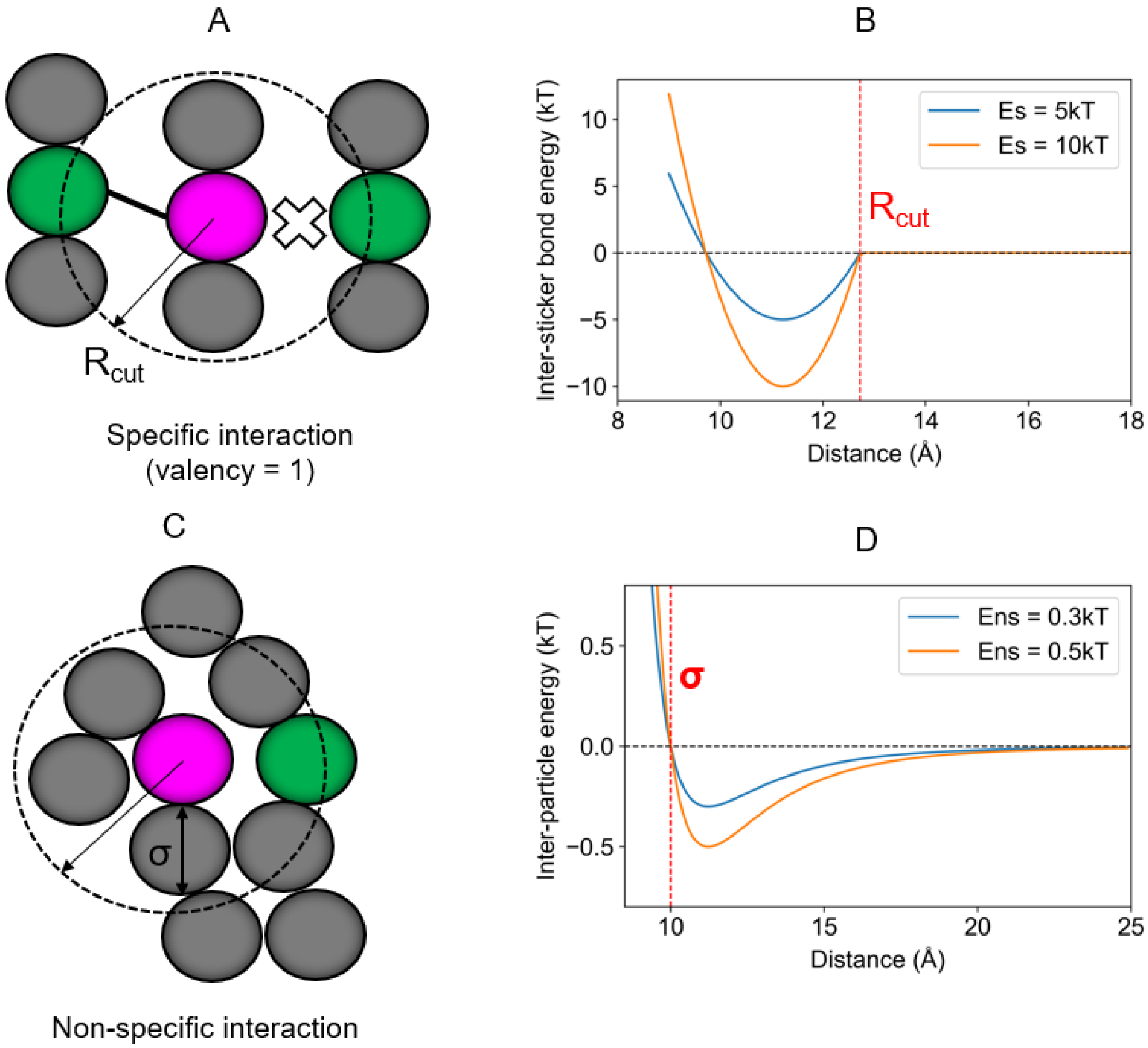


**Supplementary Figure 9: Illustration of specific and non-specific interactions.** (A) Binding sites or stickers engage in specific interactions. Once a pair of stickers (magenta and green) come closer to a cut-off distance ($R_{cut}$), they may form a "bond" with probability, $p_{on}$. The bond may dissociate with a probability, $p_{off}$. Specific interactions mimic cognate biomolecular interactions. Once a pair of stickers is bonded, they cannot engage with another sticker that may be present within $R_{cut}$. In other words, each sticker has a valency of 1. (B) The bonds are modelled with a shifted harmonic potential which becomes zero at a distance greater than $R_{cut}$. At the resting bond distance, the gain in energy is Es. In other words, the depth of energy potential is Es at the resting distance. Both probabilities ($p_{on,}$ $p_{off}$) are set to 1; hence, the stochastic factor of binding and unbinding goes away. The bond formation or breakage only depends on the inter-sticker distance. The lifetime of the bond is a function of Es, such that, $\tau_{bond} \propto e^{Es}$. The energy potential is depicted at two different Es. (C) Apart from specific interactions, each pair of beads interact via weak non-specific forces, modelled by Lennard-Jones potential. Each bead (both stickers and spacers) can exert a long-range attractive force and short-range repulsive force which determines the bead diameter (σ). One bead can interact with multiple beads, permitted by volume exclusions. The magnitude of the non-specific energy (Ens) dictates the dwelling time, that is, how much time two beads will spend on each other's vicinity. (D) Lennard-Jones potential is depicted at two different Ens.

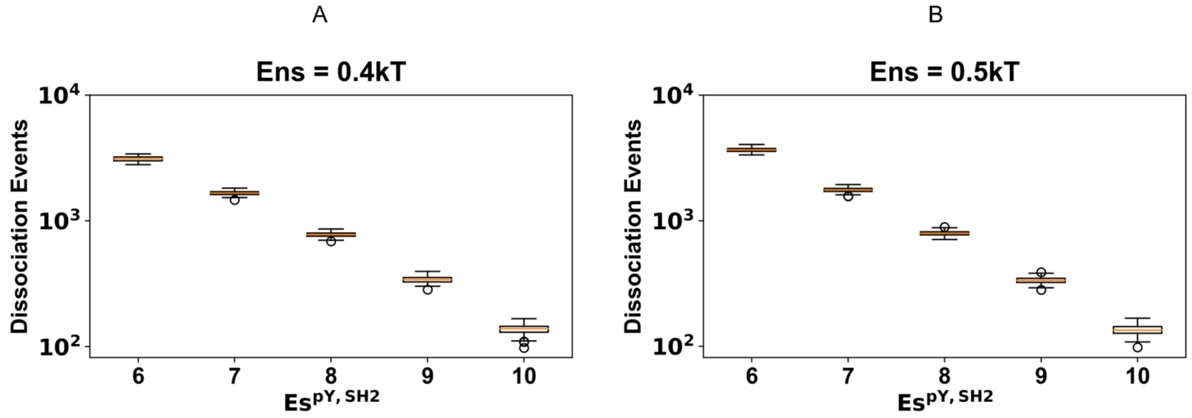


**Supplementary Figure 10: Inter-sticker dynamics.** For each type of sticker-sticker interactions, rate of dissociation, $R \propto e^{-\frac{E_S}{kT}} \Rightarrow \log\ (R) \propto -\frac{Es}{kT}$. (A, B) The number of bond dissociation events between pY and SH2 (Figure 1) at two different Ens. We note the log scale on the vertical axis. The logarithm of dissociation events vary linearly with $Es^{pY,SH2}$ with a negative slope, consistent with the rate expression that complies to detailed balance.

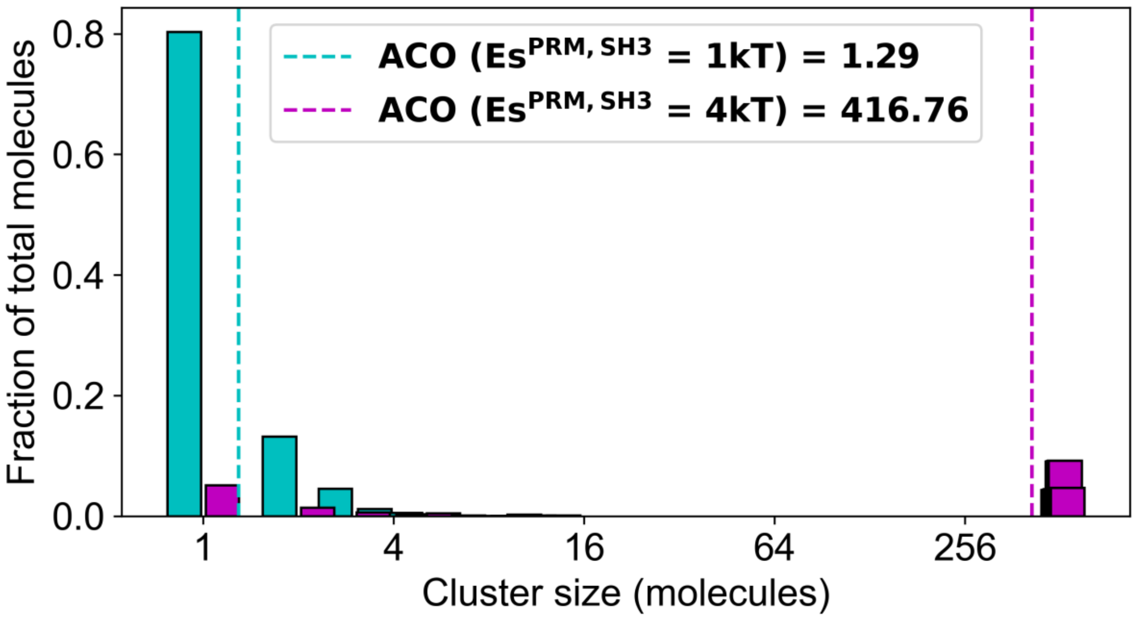


**Supplementary Figure 11: Definition of average cluster occupancy (ACO).** Cluster size distribution is quantified at two different Es. At a lower Es (cyan), the system only consists of monomers (single unconnected chains) and small oligomers. For this case, ACO = 1.29 molecules as indicated by the dashed cyan line. For a higher value of Es (magenta), large clusters emerge in the system creating a bifurcated cluster size distribution. For this case, ACO = 416.76 molecules as indicated by the dashed magenta line. ACO measures the average tendency of a molecule being part of certain cluster size. Each distribution is sampled over 20 snapshots from a relaxed trajectory.
